# Wnt5a-mediated Adipo-Cardiac Interorgan Communication in HFpEF

**DOI:** 10.1101/2025.10.29.685456

**Authors:** Anam Fatima, Faris Abusharkh, Zoe Zanetta, Sumner Gardner, Fubiao Shi, Elizabeth Kobeck, Niki Fortune, Tuerdi Subati, Katherine N. Bachmann, Jonathan D. Brown, Deepak K. Gupta, David W. J. Armstrong, Jean W. Wassenaar, James D. West, Evan L. Brittain, Sheila Collins, Anna Hemnes, Vineet Agrawal

## Abstract

**Background:** Obesity is an important risk factor in heart failure with preserved ejection fraction (HFpEF), but the precise mechanisms that drive obesity-associated cardiac remodeling are not well understood. Our previous work has shown that increased expression of the natriuretic peptide clearance receptor (*Nprc*) may drive cardiac remodeling in response to high fat diet. In this study, we hypothesized that *Nprc* plays a central role in preventing and reversing experimental HFpEF.

**Methods:** *Nprc* knockout mice were generated to induce global, cardiomyocyte-specific, or adipocyte-specific disruption of the *Nprc* (*Npr3*) gene. HFpEF was induced by the 2-hit stress of L-NAME and high fat diet. Outcomes measured included echocardiography, exercise endurance, invasive catheterization, and histologic assessment. Differential gene expression in adipocyte *Nprc* knockout visceral adipose tissue was investigated via bulk RNA sequencing and validated by RT-qPCR and culture of H9C2 cells with or without Wnt5a treatment. Adipocyte-mediated Wnt ligand release was interrupted *in vivo* via LGK974 injection in mice subjected to HFpEF conditions.

**Results:** Global inducible *Nprc* knockout prevented and reversed structural, hemodynamic, echocardiographic, and exercise tolerance features of HFpEF. This effect was found to be driven by adipocyte-specific, not cardiomyocyte-specific *Nprc* expression. Bulk RNA sequencing of adipocyte-specific *Nprc* knockout in peri-gonadal visceral adipose tissue identified downregulation of several secretory pathways, including Wnt pathways. Wnt5a was identified as one of the most downregulated genes by RNA sequencing and specifically validated by qPCR. Wnt5a exposure increased cardiomyocyte hypertrophy *in vitro*, and LGK974 (PORCN inhibitor) treatment *in vivo* decreased circulating Wnt5a levels and improved cardiac remodeling in HFpEF.

**Conclusions:** Our study identifies a novel crosstalk mechanism between cardiomyocytes and adipocytes in obesity-associated HFpEF driven by natriuretic peptide-mediated inhibition of release of Wnt5a from adipocytes.

## INTRODUCTION

Heart failure with preserved ejection fraction (HFpEF) now accounts for 50% of hospitalized heart failure patients and is associated with similar morbidity and mortality to the other forms of heart failure, yet currently has no therapies that significantly improve mortality.^1^ Despite the growing recognition of the heterogeneous nature of comorbidities that give rise to multi-system impairment in HFpEF, epidemiologic studies suggest a fundamental role for obesity in the pathogenesis of HFpEF.^2^ Obesity is independently associated with an increased risk for HFpEF relative to other forms of heart failure,^3,4^ and often presents with unique hemodynamic and cardiac structural changes relative to non-obese HFpEF.^5^ Population-wide genetic studies also suggest shared genetic risk between obesity and HFpEF, supporting a fundamental upstream role for obesity in HFpEF development.^6^

Despite these strong links, the fundamental mechanisms by which obesity selectively leads to cardiopulmonary remodeling in HFpEF is not well understood. Our prior work has identified a role for the natriuretic peptide clearance receptor in mediating obesity-mediated cardiac remodeling in model systems genetically susceptible to obesity induced HFpEF.^7,8^ Natriuretic peptides are a family of hormones secreted primarily by the heart in response to various stressors, and they exert salutatory effects in various organs through their canonical receptors (GC-A and GC-B) via particulate guanylyl cyclase-mediated generation of cGMP to counteract that stress.^9^ *Nprc* lacks a cGMP catalyzing domain and instead internalizes and degrades circulating NPs.^10^

Our prior work has identified *Nprc* as the most upregulated transcript in the heart of a genetically prone model to develop diet-induced HFpEF, but its definitive role in contributing to the cardiac pathology in HFpEF is not yet known.^7^ Based on this prior work, we hypothesized that of *Nprc* will prevent and reverse adverse cardiopulmonary remodeling in a 2-hit cardiometabolic model of HFpEF.^11^ Our findings uncovered an interorgan crosstalk mediated between adipocytes and cardiomyocytes such that natriuretic peptides inhibit the release of pro-hypertrophic proteins from adipocytes in the setting of obese HFpEF.

## METHODS

### Ethical Considerations

All animal studies were approved by the Vanderbilt University Institutional Animal Care and Use Committee (Vanderbilt University Medical Center IACUC Protocol Number M2200044, Nashville VA Medical Center IACUC Protocol Number V2200053).

### Mouse Model of Heart Failure with Preserved Ejection Fraction

All mice in the current study were bred on the C56BL6/J background. Equal numbers of 10-week-old male and female mice were used for all studies. *Npr3 ^fl/fl^* mice were generated and bred as previously described.^12^ UBC-CreERT2 (global, Jax #007001), Myh6-MerCreMer (cardiomyocyte, Jax #005657), and Adipoq-Cre (adipocyte, Jax #028020) transgenic mice were crossed with *Npr3 ^fl/fl^* to generate global and cell-specific *Nprc* knockout transgenic mice.

Inducible knockout was achieved by intraperitoneal injection of tamoxifen (2mg IP x 5) for UBC-CreERT2 mice and by 2 weeks of tamoxifen 400 mg/kg diet for cardiomyocyte specific knockout. Control groups consisted of equal numbers of Cre only or flox only mice, except for in the case of cardiomyocyte mice where Cre-only controls were used based on prior studies suggesting toxicity with Cre expression in cardiomyocytes.^13^ To induce the HFpEF phenotype as previously described,^14^ mice were first all given a calorie neutral low fat 6% lipid content diet for a 1 week run-in period (BioServ, Flemington, NJ). Thereafter, mice were randomized to either continue low fat diet or receive L-NAME in sterile water (0.5 g/L) (pH 7.0) and a 60% lipid content high fat diet ad libitum (BioServ, Flemington, NJ, USA). For LGK974 injections, LGK974 stock solutions were diluted in DMSO and a working solution with 2% DMSO and 98% corn oil was used to inject 5 mg/kg daily dose into the peritoneum via 26g x 1/2” needle daily after weight checks. Control injections consisted of 2% DMSO and 98% corn oil carrier control injections.

### Mouse Treadmill Endurance Testing

Prior to their pre-specified endpoints, mice were subjected to two days of acclimation to the treadmill at low speed of 10 m/min for 10 minutes per day. Thereafter, mice underwent a stress test protocol to identify maximal tolerated speed starting with 2 minutes of acclimation to the treadmill at 0 m/min, with subsequent increase in speed by 1 m/min every 2 minutes until exhaustion to define the maximal speed. The following day, the measured endpoint of endurance test was performed starting at 8 m/min for 10 minutes, followed by 2 m/min increase every 5 minutes until 60% of the maximal speed was obtained. Thereafter, mice were allowed to continue to walk until exhaustion, and total distance traveled was recorded.

### Mouse Echocardiography and Catheterization

Mouse echocardiography and catheterization was performed as previously reported.^7^ One day prior to catheterization, mice were anesthetized with 2-3% isoflurane. Depilatory cream was applied to the thorax. Mice were then placed on a heated table in supine position and images were acquired using the VisualSonics Vevo F2 Platform in B-mode, M-mode, and Doppler mode.

Parasternal long-axis, short-axis, and modified RV-centric views were obtained as previously described.^15^ Thereafter, mice were allowed to recover for 24 hours before undergoing anesthesia again with 2-3% isoflurane for open chest catheterization. Mice were orotracheally intubated with 22g catheter, mechanically ventilated at 18 cc/kg, and anesthetized with 2-3% vaporized isoflurane general anesthesia. On a heated surgical table in supine, ventral side-up position, a vertical incision was made along the linea alba in the rectus abdominus sheath with scissors and cautery and extended laterally. Cautery was then used to take down the diaphragm and exposure the thorax. A 1.4 French Mikro-tip catheter was directly inserted into the left ventricle for measurement of pressure and volume within the left ventricle. Afterwards, the catheter was removed and directly inserted into the right ventricle. Hemostasis of the left ventricle was ensured prior to catheterization of the right ventricle to ensure no effect of volume loss on findings. Hemodynamics were continuously recorded with a Millar MPVS-300 unit coupled to a Powerlab 8-SP analog-to-digital converter acquired at 1000 Hz and captured to a Macintosh G4 (Millar Instruments, Houston, TX). Thereafter, direct cardiac puncture was used to phlebotomize mice using a heparinized syringe and mice were sacrificed with cervical dislocation. Tissue and samples were then harvested for downstream studies from mice, including morphometric measurements.

### Cell Culture

For *in vitro* studies, H9C2 cells were cultured as previously reported.^7^ For studies investigating the effects of Wnt5a treatment, cells were transfected with either pcDNA3.2-Wnt5a plasmid (Addgene 43813) or empty vector control (Addgene 29496). All studies were conducted in H9C2 after culturing in low serum conditions (1% FBS) for 48 hours.

### Immunostaining

For hematoxylin and eosin staining, cohorts of mice were sacrificed, and hearts were cross-sectionally cut in a short-axis orientation, immediately placed in formalin for 24 hours, paraffin embedded, and sectioned (5 μm) thick for hematoxylin/eosin staining through Vanderbilt Translational Pathology Shared Resource core. For frozen sectioning, a separate cohort of hearts were 4% paraformaldehyde fixed, placed in 30% sucrose, and OCT embedded for frozen sectioning (5 μm) and wheat germ agglutinin staining. Wheat germ agglutinin staining was performed on frozen sections by permeabilizing sections (1% BSA, 0.3% TritonX, 5% goat serum) for 30 minutes, incubating slides in 100 μg/ml of Alexa Fluor 488-conjugated wheat germ agglutinin (Invitrogen, Carlsbad, CA, USA; W11261) for 1 hour in the dark, and then counterstaining with Prolong Gold Antifade with DAPI (Invitrogen, P36931) before curing for 24 hours. Imaging was performed on a Keyence BZ-X800 microscope. Quantification of cross-sectional area on WGA-stained slides was performed in a blinded fashion using the Fiji distribution of ImageJ.^16^ There were 4 individual animals used per group, and at least 3 section per animal that were quantified. For cell staining, H9C2 cells were fixed in 4% paraformaldehyde for 15 minutes on a Cellvis 33 mm culture dish (Cellvis, Mountain View, CA, USA; D35-20-1.5N) and then counterstained with ActinRed 555 (Invitrogen, R37112) and NucBlue Fixed Cell Stain (Invitrogen, R37606) according to manufacturer recommended concentrations before imaging on a Zeiss LSM 980 confocal microscope at 63x magnification. Quantification of cell size (10-20 cells per experimental replicate and 3 biologic replicates over consecutive passages 18-21) was performed in a blinded fashion using the Fiji distribution of ImageJ.^16^

### ELISA

To measure plasma levels of murine Wnt5a, 100 μl of plasma was collected and immediately flash frozen in liquid nitrogen from mice. Thereafter, the plasma was thawed and Wnt5a levels were assayed by ELISA (MyBioSource, San Diego, CA; MBS764030) following manufacturer’s included instructions and read on a BioTek Cytation 3 Plate Reader.

### RNA sequencing

Perigonadal visceral adipose tissue was harvested from mice, and approximately 50 μg of tissue was immediately placed into RLT buffer (Qiagen, Hilden, Germany). RNA isolation was then performed using the Qiagen RNeasy Plus Mini Kit (Qiagen, Hilden, Germany) following manufacturer’s guidelines and delivered to Novogene (Sacramento, California, USA) for paired-end sequencing on an Illumina platform. A nominal read depth of 20 million RNA (40 million ends) per sample were used, and actual reads ranged between 20.3 and 30.8 million reads per sample. Initial alignment and quantification were performed using the Partek Flow package.

After normalization to counts per million, 15121 genes were identified with at least one count per million and the DESeq2 package on the Partek platform was used to determine differentially expressed genes between *Nprc*-AKO and control mice. A total of 5648 genes showed at least a 50% change in expression, of which 575 genes were differentially expressed with p < 0.05 and 50% change in expression and 139 genes were differentially expressed at a more stringent p < 0.005. Gene annotation of the top 139 genes was performed using the Web Based Gene Set Analysis Toolkit (www.webgestalt.org), and gene set enrichment analysis was performed using the GSEA software (gsea-msigdb.org/gsea/index.asp).^17^

### DNA Polymerase Chain Reaction, RNA Reverse Transcription and Quantitative Polymerase Chain Reaction

For identification of sites of DNA of recombination in the Npr3 gene, DNA was isolated from tissue using the Qiagen DNeasy Blood and Tissue Kit (Qiagen) and amplified using Taq PCR Master Mix (Qiagen). For RT-qPCR, RNA was reverse transcribed to cDNA using the Qiagen One Step RT-PCR kit (Qiagen) and quantitative PCR was performed using the SYBR Green PCR master mix (Applied Biosystems, Waltham, MA, USA) on a QuantStudio 5 machine. Primer sequences used are listed in Table S1 and as previously reported.^7^

### Statistical Analysis

All statistical analysis was conducted with non-parametric testing using either Mann Whitney or Kruskal-Wallis tests in GraphPad Prism 10.

### Data Availability

RNA sequencing data will be deposited to the NCBI GEO database.

## RESULTS

### Global *Nprc* Knockout Prevents and Reverses HFpEF

Utilizing a previously generated and validated *Npr3 ^fl/fl^* mouse model,^12^ we generated an inducible global *Nprc* knockout mouse model (*Nprc*-GKO) (**Fig 1A**). After tamoxifen induction of knockout, we investigated whether global *Nprc* knockout can prevent the development of cardiopulmonary features of HFpEF in a 2-hit model of cardiometabolic stress induced by L-NAME and high fat diet for 8 weeks. Compared to control mice, global *Nprc* knockout preserved exercise endurance on a treadmill test (**Fig 1B**). Global *Nprc* knockout also prevented hypertrophy of the heart despite no significant difference in body weight (**Fig S1**), as measured by normalized heart weight (**Fig 1C**) and cardiomyocyte cross-sectional area (**Fig 1D**). On echocardiogram, global *Nprc* knockout prevented left ventricular wall hypertrophy and preserved diastolic function of the heart, measured by left atrial size and the ratio of mitral in-flow velocity (E) to mitral annular tissue Doppler velocity (e’) (**Fig 1E-G**). Finally, by invasive catheterization, global *Nprc* knockout, global *Nprc* knockout prevent an increase in left ventricular end-diastolic pressure (**Fig 1H**) and pulmonary pressure as estimated by right ventricular systolic pressure (**Fig 1I**). Global *Nprc* knockout also preserved diastolic relaxation of the right and left ventricles as measured by the relaxation constant, τ (**Fig 1J,K**). To additionally test the therapeutic potential of *Nprc* knockout, *Nprc* knockout was induced by tamoxifen treatment after 8 weeks of L-NAME and high fat diet to test the effect of *Nprc* knockout on reversing cardiopulmonary features of HFpEF (**Fig S2-A**). In this reversal model, *Nprc* knockout decreased left ventricular end-diastolic pressure (**Fig S2-C**), pulmonary pressure (**Fig S2-E**), right ventricular end-diastolic pressure (**Fig S2-F**), and biventricular diastolic relaxation (**Fig S2-D,G**) despite no statistically significant change in systemic pressure **(Fig S2-B**).

**Figure 1.**
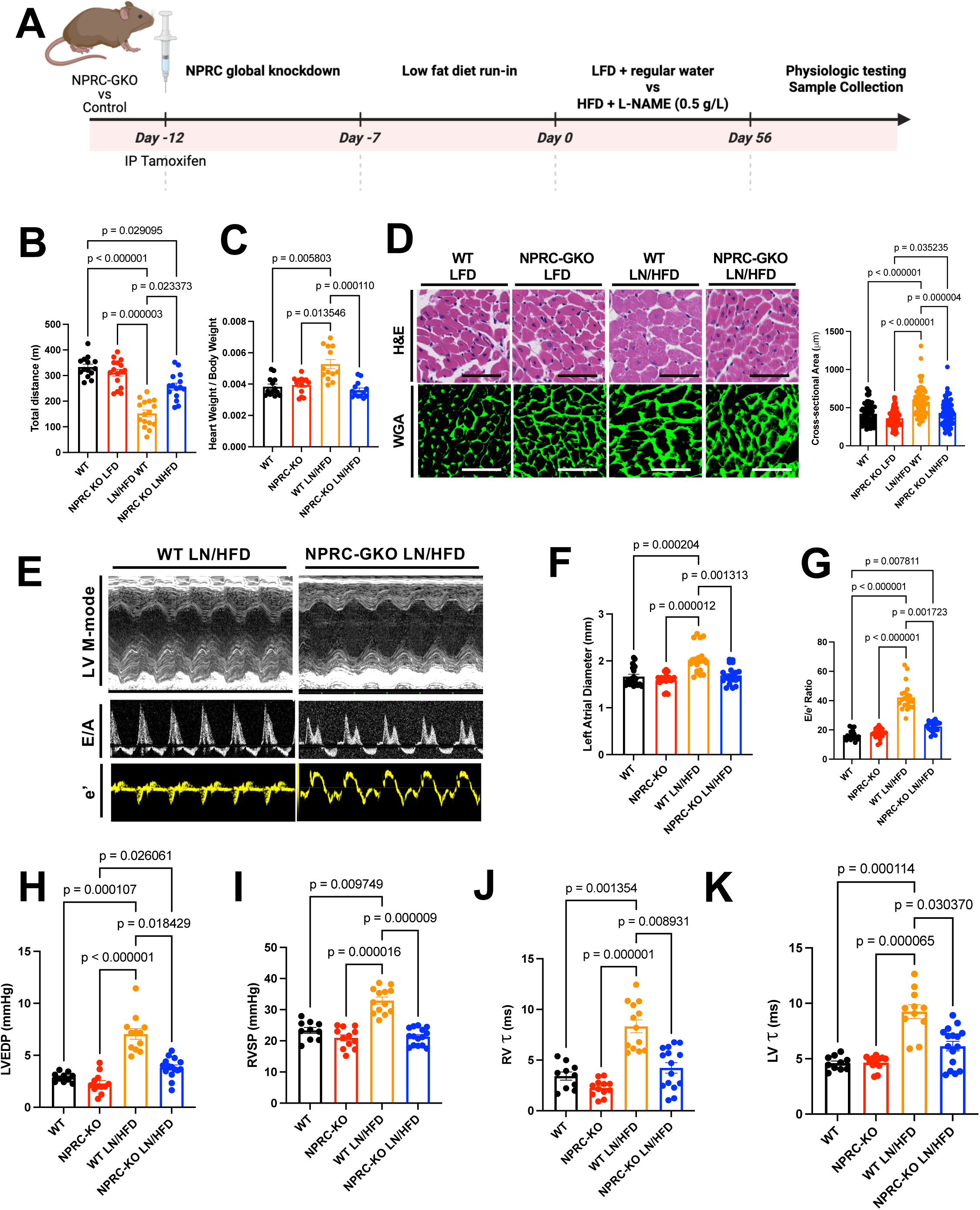
Global knockout of the natriuretic peptide clearance receptor (*Nprc*-GKO) prevents cardiac remodeling in heart failure with preserved ejection fraction (HFpEF). (A) Schematic outline of study design. (B) Total distanced traveled on treadmill in endurance test, (C) heart weight to body weight ratio, (D) histologic and wheat germ agglutinin-based hypertrophy, (E) echocardiographic assessment of left ventricle and diastolic function, (F) echocardiographic assessment of left atrial size, (G) echocardiographic assessment of diastolic measure E/e’, (H) catheter-based measurement of left ventricular end-diastolic pressure, (I) catheter-based measure of right ventricular systolic pressure, and (J-K) catheter-based measure of diastolic relaxation constant for the left and right ventricles in *Nprc*-GKO vs control mice subjected to either L-NAME and high fat diet or control water/diet for 8 weeks. *Scale bars represent 50 μm*.

### Cardiomyocyte-specific *Nprc* knockout does not protect against HFpEF

Based on prior studies that proposed that *Nprc* regulates cardiac function directly in a variety of heart failure models,^7,18^ we next hypothesized that the benefit of *Nprc* knockout is mediated by a direct effect on cardiomyocytes in the heart. To test this hypothesis, we generated cardiomyocyte-specific inducible *Nprc* knockout mice (*Nprc*-CKO) and randomized transgenic or control mice to HFpEF inducing or control conditions for 8 weeks (**Fig 2A**). Cardiomyocyte-specific knockout of the *Npr3* gene was identified by recombination of exon 3 in the *Npr3* gene in the heart, and by RT-qPCR of *Npr3* expression (**Fig S3**). Cardiomyocyte-specific *Nprc* knockout did not prevent weight gain (**Fig S1**), cardiomyocyte hypertrophy (**Fig 2B,D**), exercise tolerance (**Fig 2C**), end-systolic or end-diastolic pressure in the left or right ventricle (**Fig 2E,G-I**), or diastolic relaxation of the right or left ventricle (**Fig 2F,J**). These findings suggested that the protective effect of *Nprc* knockout was not mediated by a direct effect on cardiomyocytes.

**Figure 2.**
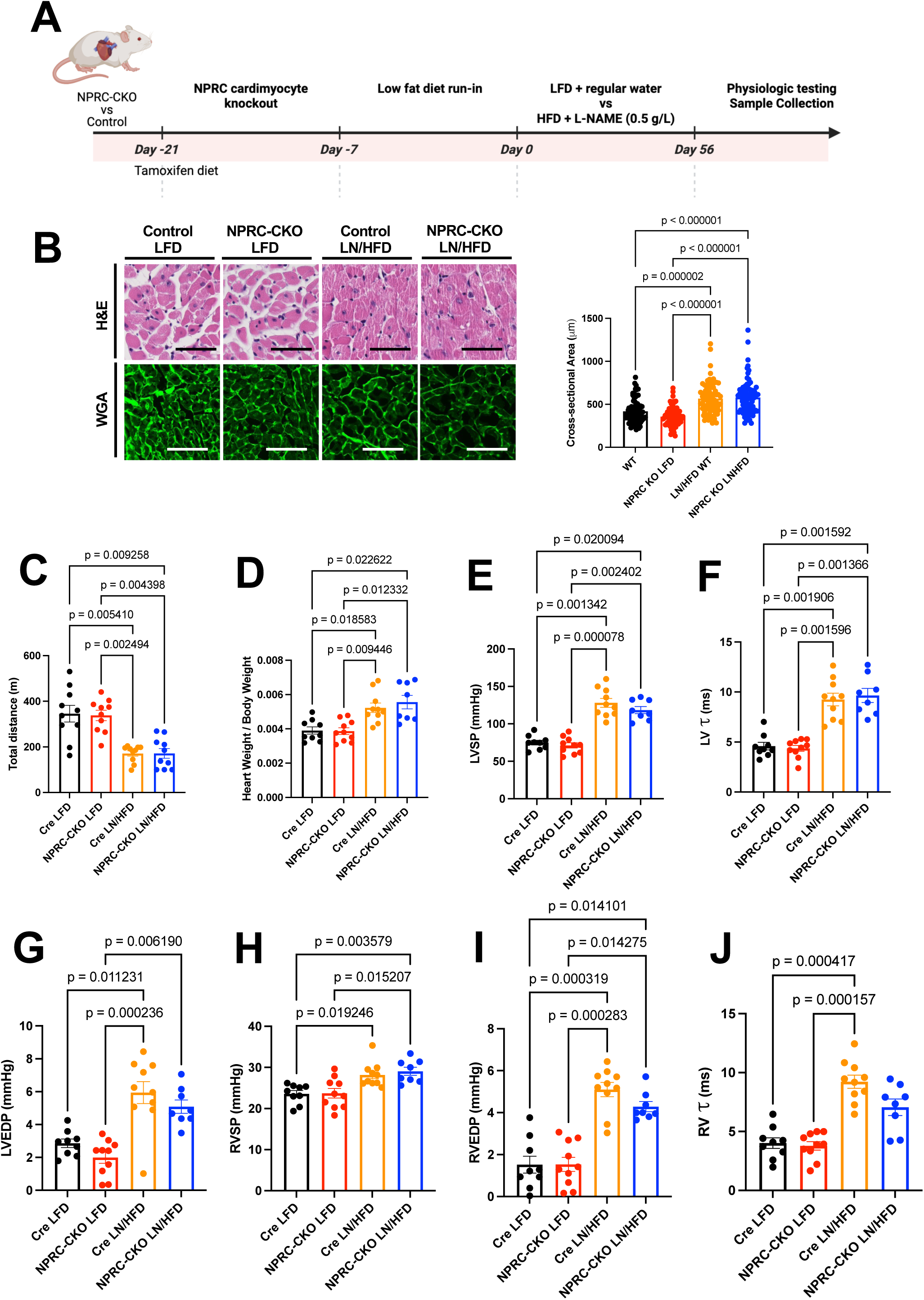
Cardiomyocyte-specific knockout of the natriuretic peptide clearance receptor (*Nprc*-CKO) does not prevent cardiac remodeling in heart failure with preserved ejection fraction (HFpEF). (A) Schematic outline of study design. (B) Histologic and wheat germ agglutinin-based hypertrophy, (C) total distanced traveled on treadmill in endurance test, (D) heart weight to body weight ratio, (E) left ventricular systolic pressure, (F) diastolic relaxation of the left ventricle, (G) left ventricular end-diastolic pressure, (H) right ventricular systolic pressure, (I) right ventricular end-diastolic pressure, and (J) diastolic relaxation of the right ventricle in *Nprc*-CKO vs control mice subjected to either L-NAME and high fat diet or control water/diet for 8 weeks. *Scale bars represent 50 μm*.

### Adipocyte-specific *Nprc* knockout prevents cardiac remodeling in HFpEF

Based on prior literature, NRPC is known to be highly expressed in adipocytes, and dynamically regulated by both environmental and nutritional changes.^19^ Mice lacking *Nprc* in adipocytes, despite only mild changes in weight gain with diet-induced obesogenic conditions, display marked improvements in insulin resistance and glucose tolerance with an associated activation of the thermogenic program in adipocytes.^12^ Based on this literature, we hypothesized that the protective effect of *Nprc* knockout might be mediated an interorgan communication between adipose tissue and the heart. To test this hypothesis, we utilized previously generated adipocyte-specific *Nprc* knockout mice^12^ and subjected mice or control counterparts to HFpEF-inducing conditions for 8 weeks (**Fig 3A**). Mice lacking *Nprc* expression in adipocytes displayed preserved cardiomyocyte size (**Fig 3B**), left ventricular dimensions and diastolic function (**Fig 3C,D,F-G**) by echocardiography, and exercise endurance (**Fig 3E**) despite no significant change in weight gain during the experimental period (**Fig S1**). By invasive catheterization, despite no differences in end-systolic pressure of the left ventricle (**Fig 3H**) or the right ventricle (**Fig 3K**), mice lack adipocyte-specific *Nprc* expression displayed preserved end-diastolic pressure and diastolic relaxation of both ventricles (**Fig 3I-J,L-M**), suggesting that adipocyte-specific *Nprc* knockout improved the heart’s response to a sustained load stress in HFpEF.

**Figure 3.**
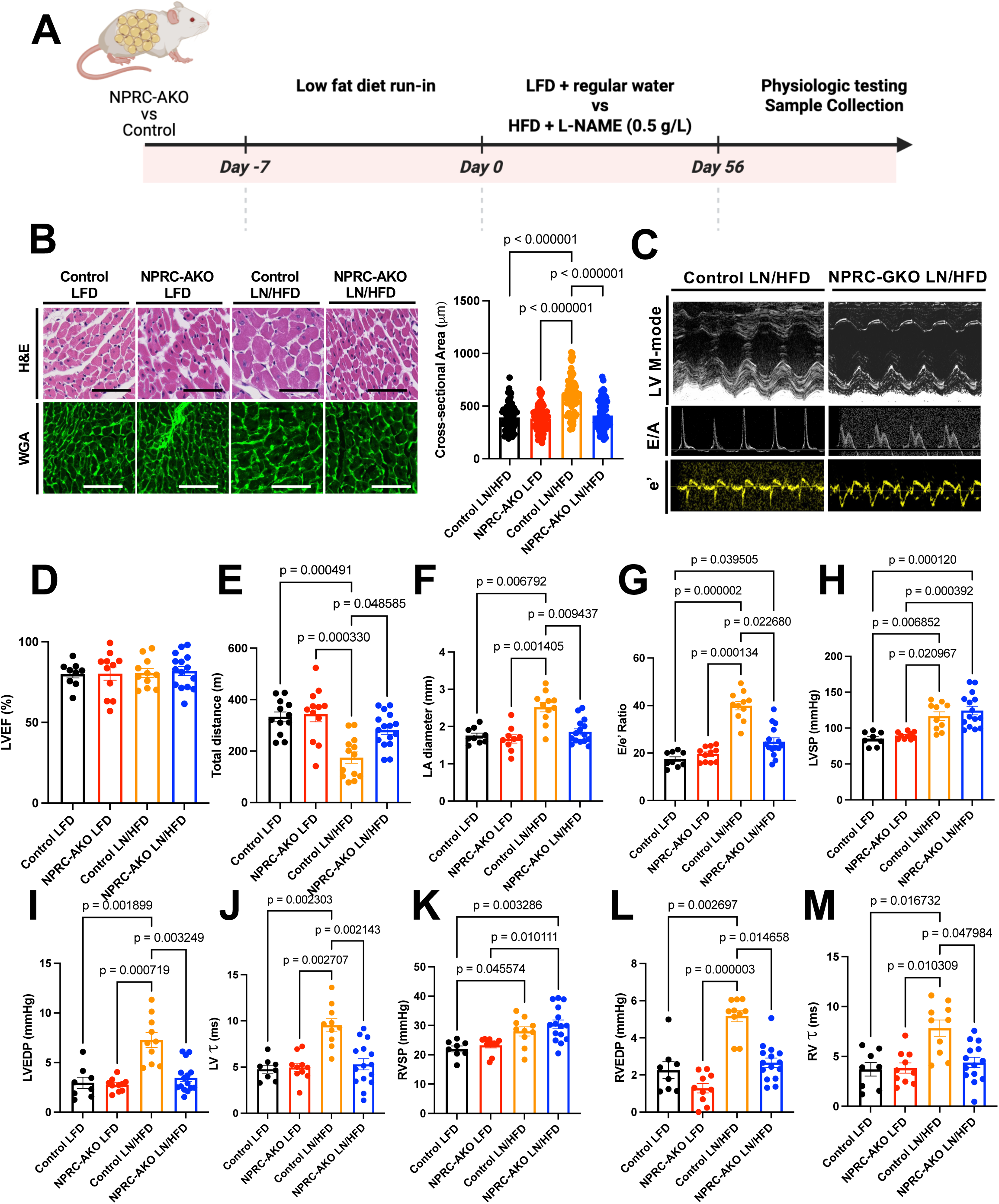
Adipocyte-specific knockout of the natriuretic peptide clearance receptor (*Nprc*-AKO) prevents cardiac remodeling in heart failure with preserved ejection fraction (HFpEF). (A) Schematic outline of study design. (B) Histologic and wheat germ agglutinin-based hypertrophy, (C) echocardiographic assessment of the left ventricle systolic and diastolic function, (D) left ventricular ejection fraction (%), (E) total distanced traveled on treadmill in endurance test, (F) left atrial size by echo, (G) E/e’ ratio by echo, (H) left ventricular systolic pressure, (I) left ventricular end-diastolic pressure, (J) diastolic relaxation of the left ventricle, (K) right ventricular systolic pressure, (L) right ventricular end-diastolic pressure, and (M) diastolic relaxation of the right ventricle in *Nprc*-AKO vs control mice subjected to either L-NAME and high fat diet or control water/diet for 8 weeks. *Scale bars represent 50 μm*.

### *Nprc* knockout in Adipocytes Alters the Adipose Secretome

To understand how *Nprc* knockout in adipocytes may regulate cardiac remodeling in HFpEF, we next isolated perigonadal visceral adipose tissue from *Nprc*-AKO or control mice after 8 weeks of HFpEF-inducing conditions and used bulk RNA sequencing to identify differentially expressed genes (**Fig 4A**). Adipocyte-specific knockout has previously been confirmed.^12^ Among 15,121 genes with at least one count per million transcript level expression per sample, a total of 5648 genes showed at least a 50% change in expression. Among those genes, 575 were differentially expressed at a nominal, unadjusted p-value of 0.05, and a total of 139 genes were differentially expressed at a p-value < 0.01. Gene enrichment gene ontology of the 139 genes revealed that 42 out of the 139 genes (30%) encoded for proteins that were known to be secreted (**Fig 4B**). Gene set enrichment analysis identified a number of pathways that were upregulated in *Nprc*-AKO adipose tissue including a number of metabolic pathways involving mitochondrial respiratory chain function, oxidative phosphorylation, mTORC1 signaling, and insulin signaling (**Fig 4C**), all pathways that were consistent with previous studies supporting the metabolic benefits of adipocyte-specific *Nprc* knockout.^12,19^ However, gene set enrichment analysis also identified a number of downregulated pathways involving secretory protein-related signaling pathways related to the epidermal growth factor family, fibroblast growth factor and receptor family, and Wnt family (**Fig 4C**). Among the top 139 differentially expressed genes, Wnt5a and Wnt9a were significantly downregulated in expression in *Nprc*-AKO adipose tissue (**Fig 4D**).

**Figure 4.**
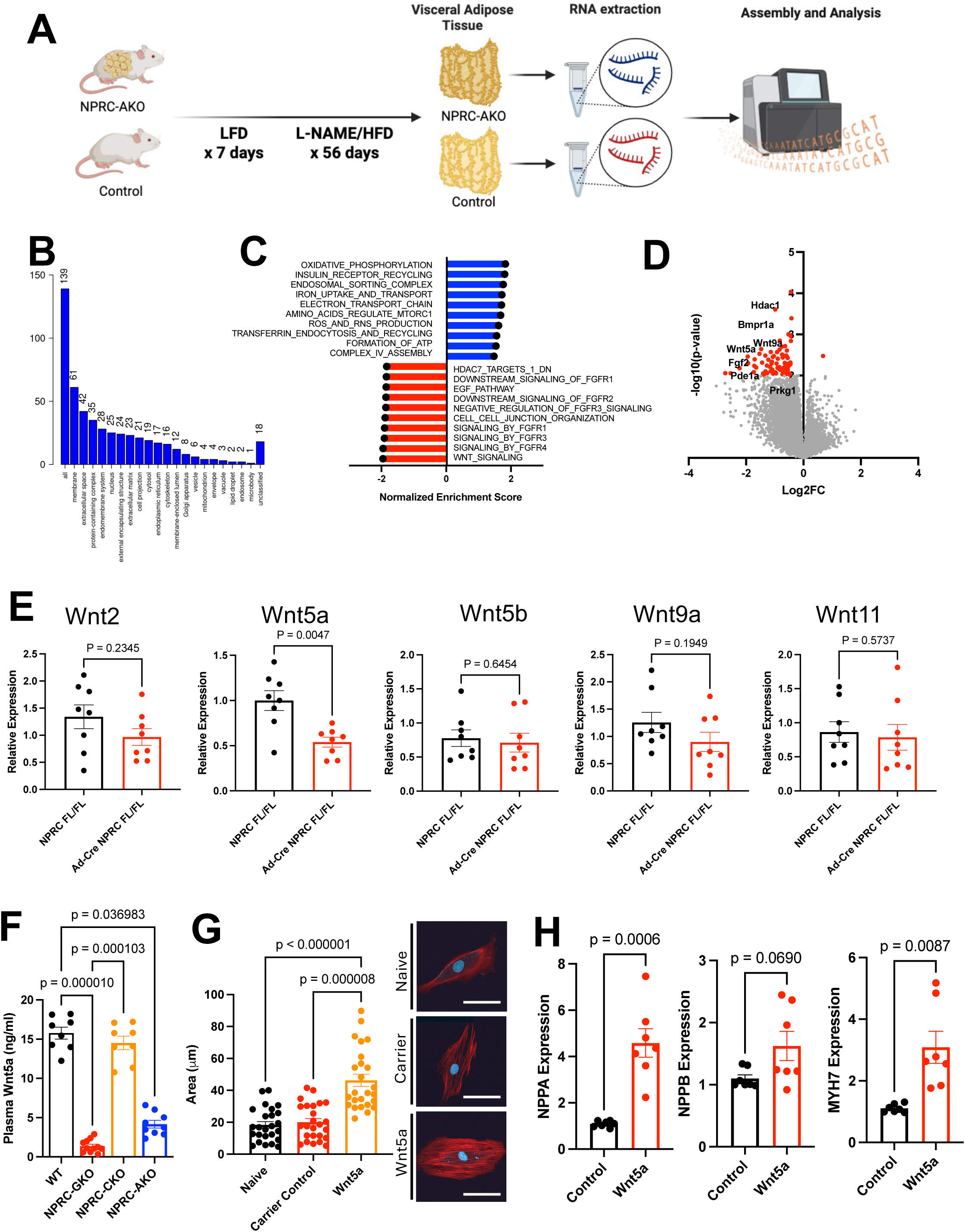
Adipocyte-specific *Nprc* knockout (*Nprc*-AKO) in adipocytes decreases Wnt5a expression. (A) Schematic outline of study. (B) Cellular localization of differentially expressed genes (DEGs) in *Nprc*-AKO visceral adipose tissue (VAT). (C) Increased and decreased gene ontologies in *Nprc*-AKO vs control VAT. (D) Volcano plot of DEGs in *Nprc*-AKO vs control VAT (red indicates p < 0.01 and 50%-fold-change compared to control). (E) Reverse transcriptase quantitative PCR of top 5 Wnt isoforms in *Nprc*-AKO vs control VAT. (F) Plasma circulating levels of Wnt5a in global (GKO), cardiomyocyte-specific (CKO), adipocyte-specific (AKO), and control (WT) mice after L-NAME and high fat diet. (G) Cell size in H9C2 cells after treatment with Wnt5a. (H) RT-qPCR for hypertrophic markers NPPA, NPPB, and MYH7 in H9C2 cells after Wnt5a treatment.

Given that the Wnt signaling pathway was the most downregulated pathway by gene set enrichment analysis, we sought to validate the top 5 differentially expressed Wnt isoforms identified by bulk RNA sequencing using RT-qPCR. Among the top 5 Wnt isoforms, only Wnt5a was significantly decreased in a second cohort of *Nprc*-AKO vs control mice after 8 weeks of HFpEF-inducing conditions (**Fig 4E**). Based on studies suggesting circulating Wnt5a levels are elevated in patients with heart failure,^20,21^ we next asked whether Wnt5a expression in adipose tissue correlates with circulating levels of Wnt5a in our murine model system. We found that Wnt5a levels were elevated in wild type and *Nprc*-CKO mice after 8 weeks of HFpEF-inducing conditions (**Fig 4F**), whereas Wnt5a levels were low and near normal levels in *Nprc*-GKO and *Nprc*-AKO mice (**Fig 4F**).^22^ Finally, we asked the question of whether Wnt5a was sufficient to activate a hypertrophic gene program in cardiomyocytes. To address this question, we investigated cell size and expression of a hypertrophic gene program in H9C2 cells treated with either Wnt5a or empty vector plasmid. After 48 hours of expression, we found that Wnt5a-treated cells were increased in size (**Fig 4G**) and activated a hypertrophic gene program characterized by increased expression of *Nppa*, *Nppb*, and *Myh7* (**Fig 4H**).

### Interruption of Adipocyte-Cardiomyocyte Crosstalk Mitigates Cardiac Remodeling in HFpEF

To investigate whether interruption of adipocyte-mediated Wnt5a may alleviate cardiopulmonary remodeling, we next treated wild type C57BL/6J mice subjected to 8 weeks of HFpEF-inducing conditions with intraperitoneal injections of LGK974 or carrier control for 4 weeks (**Fig 5A**).

**Figure 5.**
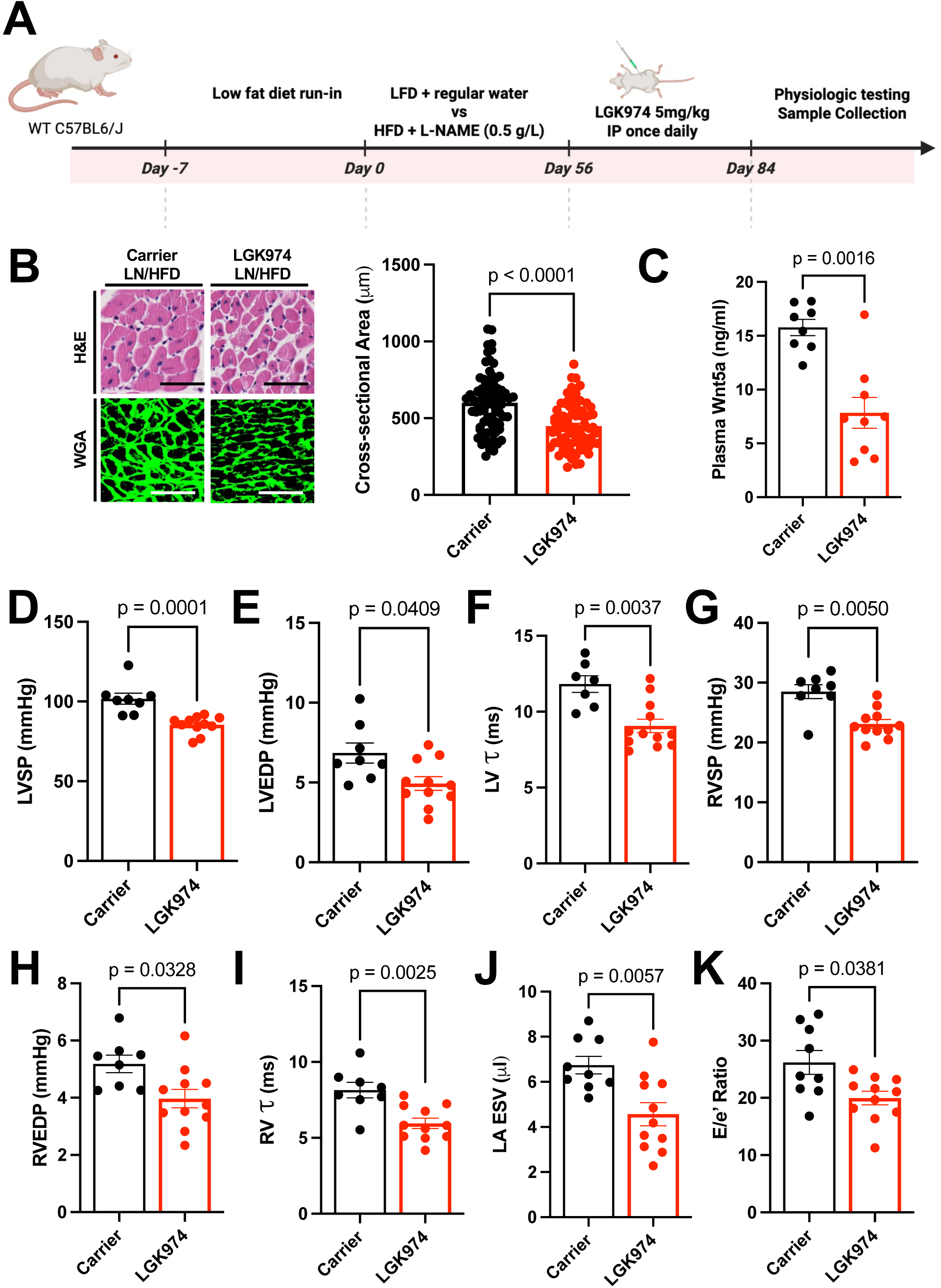
Inhibition of Wnt release by treatment with LGK974 mitigates cardiac remodeling in HFpEF. (A) Schematic outline of study. (B) Histologic and wheat germ agglutinin-based hypertrophy, (C) plasma Wnt5a levels, (D) left ventricular systolic pressure, (E) left ventricular end-diastolic pressure, (F) diastolic relaxation of the left ventricle, (G) right ventricular systolic pressure, (H) right ventricular end-diastolic pressure, (I) diastolic relaxation of the right ventricle, (J) left atrial end-systolic volume by echocardiography, and (K) E/e’ ratio by echocardiography in C57/BL6J mice treated with 8 weeks of L-NAME and high fat diet, followed by LGK974 or carrier control intraperitoneal injections.

**Figure 6.**
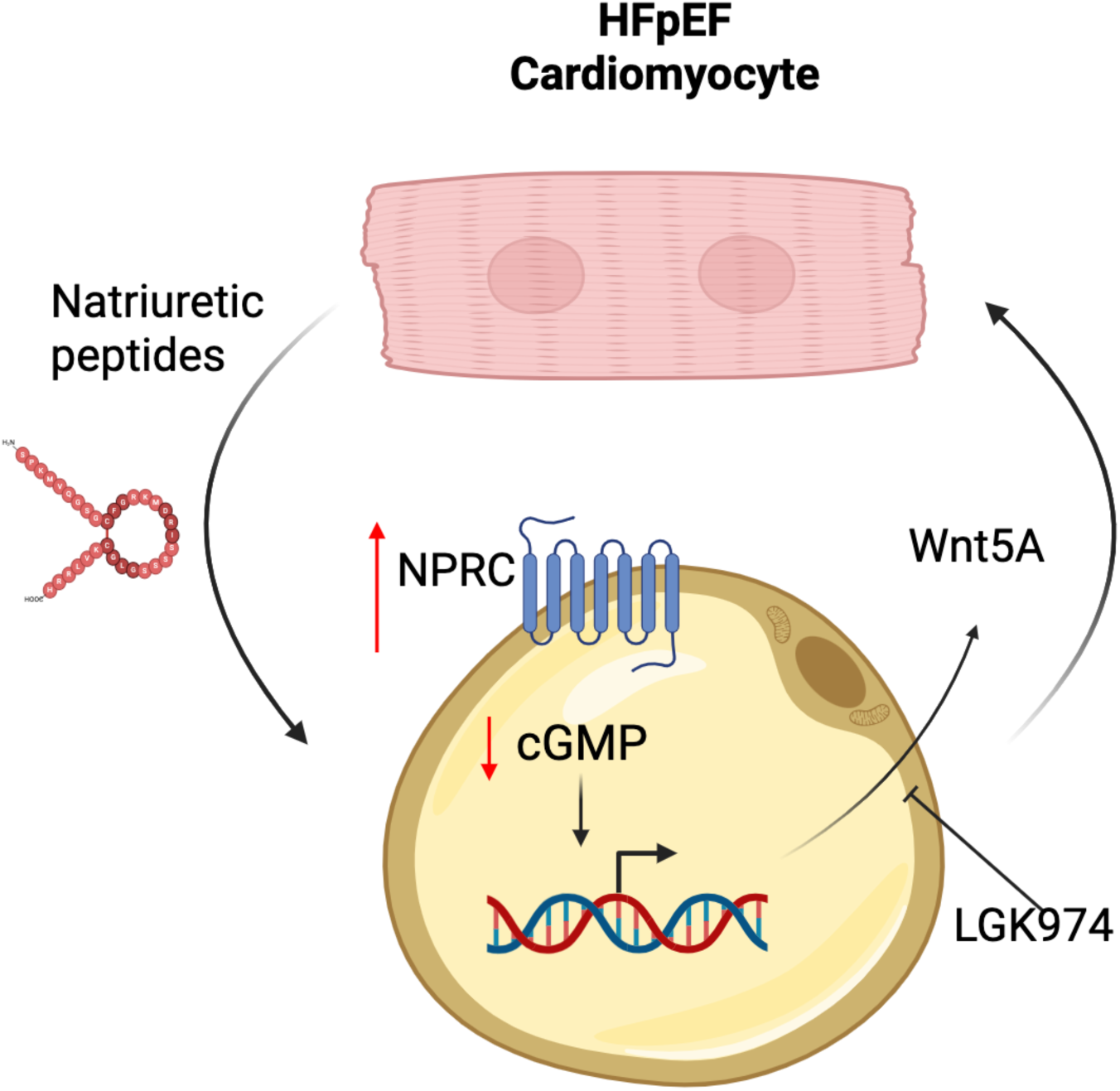
Working model for interorgan crosstalk between cardiomyocytes and adipocytes in obesity-associated HFpEF.

LGK974 is an inhibitor of the acyltransferase, porcupine, which is required for palmitoylation of Wnt proteins and subsequent secretion and signaling of Wnt proteins.^23,24^ LGK974 has been studied in Phase I and II clinical trials as a cancer therapeutic.^25,26^ We first confirmed that intraperitoneal injection of LGK974 only affected visceral organs by measuring transcriptional targets of Wnt signaling after a single injection of LGK974 (**Fig S4**). We next tested the therapeutic effect of LGK974 in HFpEF by treating mice with 4 weeks of LGK974 after 8 weeks of HFpEF induction (**Fig 5A**). Compared to control injection, LGK974 treatment led to a decrease in circulating Wnt5a plasma levels (**Fig 5C**) as well as a decrease in cardiomyocyte hypertrophy as measured by cross-sectional area (**Fig 5B**). Invasive catheterization demonstrated an improvement in end-systolic pressure, end-diastolic pressure, diastolic relaxation of the right and left ventricle (**Fig 5D-I**), and echocardiographic assessment demonstrated an improvement in left atrial end-systolic volume and diastolic function as assessed by E/e’ (**Fig 5J,K**).

## DISCUSSION

The overarching goal of this study was to better understand mechanisms that may drive adverse cardiopulmonary remodeling in the context of obesity-associated heart failure with preserved ejection fraction (HFpEF). Based on our prior work and literature, we hypothesized that restoration of natriuretic peptide signaling in obese HFpEF would prevent and reverse cardiac remodeling in HFpEF. The findings of our study demonstrate that the protective effect of natriuretic peptide signaling, restored by knockout of the natriuretic peptide clearance receptor (*Nprc*), is driven by an adipocyte-specific regulation of Wnt5a release.

Despite recognition of HFpEF as a highly heterogeneous syndrome driven by varying upstream comorbidities, obesity and diabetes have consistently been identified as features of a unique endophenotype of HFpEF in unbiased approaches. ^27–31^ Based on these observations as well as the recognition of increasing prevalence of obesity in more contemporary HFpEF cohorts,^2^ numerous clinical studies have now demonstrated symptomatic improvement from targeting obesity and diabetes through lifestyle, surgical, and pharmacologic interventions modifications.^32–38^ A number of studies, however, have also suggested that the link between obesity and HFpEF is more complex than just weight alone. As with other aspects of obesity comorbidities, location of adiposity has been shown to strongly influence outcomes in HFpEF, and the symptomatic benefit from weight-targeting therapies appear to be at least partially independent of the degree of weight loss.^5,39,40^

Pre-clinical and translational studies have also suggested that underlying genetic background may influence the obesity-associated risk of adverse remodeling,^7,41,42^ collectively suggesting a more complex interplay between adiposity and cardiac remodeling in HFpEF that requires further mechanistic understanding.

Our study contributes to the growing understanding that adipose tissue is a highly active endocrine organ that strongly contributes to circulating hormones that may be biologically active in cardiovascular disease.^43,44^ Among other secretory pathways such as the EGF, FGF, and Wnt secretory pathways regulated by downregulation of the natriuretic peptide clearance receptor in our study, we identified and further validated an important role for Wnt5a in mediating obesity-associated cardiac remodeling. This finding is supported by human studies that identified selective increases in Wnt5a levels in the visceral and epicardial adipose tissue of patients as opposed to subcutaneous fat,^45,46^ consistent with the depots of adipose tissue that are typically associated with HFpEF risk and mortality.^39,47,48^ Furthermore, studies have not only found that circulating levels of WNT5A are predictive of outcomes in patients with heart failure, but that the non-canonical receptor for Wnt5a, ROR2, is one of the most upregulated genes in the setting of heart failure.^20,21,46,49^ Finally, other studies, including small and large animal models of HFpEF, have similarly demonstrated benefit from pharmacologic inhibition of Wnt signaling.^50,51^ Our study contributes to this existing literature by identifying the visceral adipose tissue as an important source of these Wnt ligands in the context of obesity-associated HFpEF, and by identifying the upstream role of natriuretic peptide signaling in modulating Wnt release from adipose tissue.

Natriuretic peptide signaling has long been established as a key pathway that is dysregulated in all forms of heart failure.^52^ Multiple mechanisms contribute to deficient natriuretic peptide signaling in heart failure, including deficient production of natriuretic peptides, increased enzymatic degradation by neprilysin, and impaired downstream signaling by PDE-9.^53–55^

Notably, while previous studies have suggestive *Nprc* plays a protective role in direct myocardial injury as a result of ischemic insult,^18^ our study suggests that *Nprc* may play a more deleterious role in metabolically driven cardiac remodeling, highlighting the complex and cell-specific dynamics of natriuretic peptide signaling in disease. Our study contributes to a growing body of literature that also implicates a deleterious role for *Nprc* in mediating end-organ damage from metabolic syndrome,^7,12,19,56–58^ Specifically, prior work has shown that cGMP signaling is amplified after adipocyte-specific knockout of *Nprc* and drives activation of a *UCP1*-mediated thermogenic program towards a beige/brown fat phenotype, which may be an important driver of systemic protection.^7,12,19^ Our study indicates *Nprc* regulates the adipocyte secretome, providing a novel mechanism by which natriuretic peptides exert anti-remodeling actions in the heart.

Our study has several limitations. HFpEF is well established to be a heterogeneous syndrome driven by varying upstream risk factors. This study was conducted in one model of obesity-associated HFpEF driven by the 2-hit stress of L-NAME and high fat diet. While Wnt inhibition has been shown to be beneficial in the context of other non-obese models of HFpEF such salt-sensitive hypertension models (DOCA+salt),^50,51^ more work is necessary to define the precise mechanisms that drive interorgan and Wnt-mediated crosstalk in other phenotypes of HFpEF. Our study also specifically focused on the role of Wnt signaling in mediating adipocyte-cardiomyocyte crosstalk, but we acknowledge that the adipocyte secretome is complex and likely includes many secreted proteins and metabolites. Further work is necessary to fully define the adipocyte secretome in HFpEF, dissect out the various contributions of different secreted products, and understand the extent to which different depots of adipose tissue may differentially affect cardiac remodeling.

In summary, our current study identifies an important axis of crosstalk that occurs between cardiomyocytes and adipocytes in the setting of obesity-associated HFpEF where *Nprc* expression in adipocytes regulates adipocyte-cardiac myocyte crosstalk by attenuating the release of pro-hypertrophic Wnt5a from adipocytes. These data support the hypothesis that adipokine signaling plays a key role in the pathogenesis of HFpEF,^59^ which may be partially mediated by deficient NP signaling.

## REFERENCES CITED

1. Shah KS, Xu H, Matsouaka RA, Bhatt DL, Heidenreich PA, Hernandez AF, Devore AD, Yancy CW, Fonarow GC. Heart Failure With Preserved, Borderline, and Reduced Ejection Fraction: 5-Year Outcomes. J Am Coll Cardiol. 2017;70:2476–2486. doi: 10.1016/j.jacc.2017.08.074

2. Rao VN, Fudim M, Mentz RJ, Michos ED, Felker GM. Regional adiposity and heart failure with preserved ejection fraction. Eur J Heart Fail. 2020;22:1540–1550. doi: 10.1002/ejhf.1956

3. Savji N, Meijers WC, Bartz TM, Bhambhani V, Cushman M, Nayor M, Kizer JR, Sarma A, Blaha MJ, Gansevoort RT, et al. The Association of Obesity and Cardiometabolic Traits With Incident HFpEF and HFrEF. JACC Heart Fail. 2018;6:701–709. doi: 10.1016/j.jchf.2018.05.018

4. Rao VN, Zhao D, Allison MA, Guallar E, Sharma K, Criqui MH, Cushman M, Blumenthal RS, Michos ED. Adiposity and Incident Heart Failure and its Subtypes: MESA (Multi-Ethnic Study of Atherosclerosis). JACC Heart Fail. 2018;6:999–1007. doi: 10.1016/j.jchf.2018.07.009

5. Obokata M, Reddy YNV, Pislaru SV, Melenovsky V, Borlaug BA. Evidence Supporting the Existence of a Distinct Obese Phenotype of Heart Failure With Preserved Ejection Fraction. Circulation. 2017;136:6–19. doi: 10.1161/CIRCULATIONAHA.116.026807

6. Joseph J, Liu C, Hui Q, Aragam K, Wang ZY, Charest B, Huffman JE, Keaton JM, Edwards TL, Demissie S, et al. Genetic architecture of heart failure with preserved versus reduced ejection fraction. Nature Communications. 2022;13. doi: ARTN 7753 10.1038/s41467-022-35323-0

7. Agrawal V, Fortune N, Yu S, Fuentes J, Shi F, Nichols D, Gleaves L, Poovey E, Wang TJ, Brittain EL, et al. Natriuretic peptide receptor C contributes to disproportionate right ventricular hypertrophy in a rodent model of obesity-induced heart failure with preserved ejection fraction with pulmonary hypertension. Pulm Circ. 2019;9:2045894019878599. doi: 10.1177/2045894019895452

8. Meng Q, Lai YC, Kelly NJ, Bueno M, Baust JJ, Bachman TN, Goncharov D, Vanderpool RR, Radder JE, Hu J, et al. Development of a Mouse Model of Metabolic Syndrome, Pulmonary Hypertension, and Heart Failure with Preserved Ejection Fraction. Am J Respir Cell Mol Biol. 2017;56:497–505. doi: 10.1165/rcmb.2016-0177OC

9. Lee CY, Burnett JC, Jr. Natriuretic peptides and therapeutic applications. Heart Fail Rev. 2007;12:131–142. doi: 10.1007/s10741-007-9016-3

10. Matsukawa N, Grzesik WJ, Takahashi N, Pandey KN, Pang S, Yamauchi M, Smithies O. The natriuretic peptide clearance receptor locally modulates the physiological effects of the natriuretic peptide system. Proc Natl Acad Sci U S A. 1999;96:7403– 7408. doi: 10.1073/pnas.96.13.7403

11. Schiattarella GG, Altamirano F, Tong D, French KM, Villalobos E, Kim SY, Luo X, Jiang N, May HI, Wang ZV, et al. Nitrosative stress drives heart failure with preserved ejection fraction. Nature. 2019;568:351–356. doi: 10.1038/s41586-019-1100-z

12. Wu W, Shi F, Liu D, Ceddia RP, Gaffin R, Wei W, Fang H, Lewandowski ED, Collins S. Enhancing natriuretic peptide signaling in adipose tissue, but not in muscle, protects against diet-induced obesity and insulin resistance. Sci Signal. 2017;10. doi: 10.1126/scisignal.aam6870

13. Koitabashi N, Bedja D, Zaiman AL, Pinto YM, Zhang M, Gabrielson KL, Takimoto E, Kass DA. Avoidance of transient cardiomyopathy in cardiomyocyte-targeted tamoxifen-induced MerCreMer gene deletion models. Circ Res. 2009;105:12–15. doi: 10.1161/CIRCRESAHA.109.198416

14. Agrawal V, Kropski JA, Gokey JJ, Kobeck E, Murphy MB, Murray KT, Fortune NL, Moore CS, Meoli DF, Monahan K, et al. Myeloid Cell Derived IL1beta Contributes to Pulmonary Hypertension in HFpEF. Circ Res. 2023;133:885–898. doi: 10.1161/CIRCRESAHA.123.323119

15. Brittain E, Penner NL, West J, Hemnes A. Echocardiographic assessment of the right heart in mice. J Vis Exp. 2013. doi: 10.3791/50912

16. Schindelin J, Arganda-Carreras I, Frise E, Kaynig V, Longair M, Pietzsch T, Preibisch S, Rueden C, Saalfeld S, Schmid B, et al. Fiji: an open-source platform for biological-image analysis. Nat Methods. 2012;9:676–682. doi: 10.1038/nmeth.2019

17. Subramanian A, Tamayo P, Mootha VK, Mukherjee S, Ebert BL, Gillette MA, Paulovich A, Pomeroy SL, Golub TR, Lander ES, et al. Gene set enrichment analysis: a knowledge-based approach for interpreting genome-wide expression profiles. Proc Natl Acad Sci U S A. 2005;102:15545–15550. doi: 10.1073/pnas.0506580102

18. Lowe VJ, Aubdool AA, Moyes AJ, Dignam JP, Perez-Ternero C, Baliga RS, Smart N, Hobbs AJ. Cardiomyocyte-derived C-type natriuretic peptide diminishes myocardial ischaemic injury by promoting revascularisation and limiting fibrotic burden. Pharmacol Res. 2024;209:107447. doi: 10.1016/j.phrs.2024.107447

19. Shi F, Simandi Z, Nagy L, Collins S. Diet-dependent natriuretic peptide receptor C expression in adipose tissue is mediated by PPARgamma via long-range distal enhancers. J Biol Chem. 2021;297:100941. doi: 10.1016/j.jbc.2021.100941

20. Abraityte A, Lunde IG, Askevold ET, Michelsen AE, Christensen G, Aukrust P, Yndestad A, Fiane A, Andreassen A, Aakhus S, et al. Wnt5a is associated with right ventricular dysfunction and adverse outcome in dilated cardiomyopathy. Sci Rep. 2017;7:3490. doi: 10.1038/s41598-017-03625-9

21. Abraityte A, Vinge LE, Askevold ET, Lekva T, Michelsen AE, Ranheim T, Alfsnes K, Fiane A, Aakhus S, Lunde IG, et al. Wnt5a is elevated in heart failure and affects cardiac fibroblast function. J Mol Med (Berl). 2017;95:767–777. doi: 10.1007/s00109-017-1529-1

22. Chi S, Xue J, Chen X, Liu X, Ji Y. Correlation of plasma and urine Wnt5A with the disease activity and cutaneous lesion severity in patients with systemic lupus erythematosus. Immunol Res. 2022;70:174–184. doi: 10.1007/s12026-021-09253-w

23. Lung H, Wentworth KL, Moody T, Zamarioli A, Ram A, Ganesh G, Kang M, Ho S, Hsiao EC. Wnt pathway inhibition with the porcupine inhibitor LGK974 decreases trabecular bone but not fibrosis in a murine model with fibrotic bone. JBMR Plus. 2024;8:ziae011. doi: 10.1093/jbmrpl/ziae011

24. Liu J, Pan S, Hsieh MH, Ng N, Sun F, Wang T, Kasibhatla S, Schuller AG, Li AG, Cheng D, et al. Targeting Wnt-driven cancer through the inhibition of Porcupine by LGK974. Proc Natl Acad Sci U S A. 2013;110:20224–20229. doi: 10.1073/pnas.1314239110

25. Tabernero J, Van Cutsem E, Garralda E, Tai D, De Braud F, Geva R, van Bussel MTJ, Fiorella Dotti K, Elez E, de Miguel MJ, et al. A Phase Ib/II Study of WNT974 + Encorafenib + Cetuximab in Patients With BRAF V600E-Mutant KRAS Wild-Type Metastatic Colorectal Cancer. Oncologist. 2023;28:230–238. doi: 10.1093/oncolo/oyad007

26. Rodon J, Argiles G, Connolly RM, Vaishampayan U, de Jonge M, Garralda E, Giannakis M, Smith DC, Dobson JR, McLaughlin ME, et al. Phase 1 study of single-agent WNT974, a first-in-class Porcupine inhibitor, in patients with advanced solid tumours. Br J Cancer. 2021;125:28–37. doi: 10.1038/s41416-021-01389-8

27. Shah SJ, Katz DH, Selvaraj S, Burke MA, Yancy CW, Gheorghiade M, Bonow RO, Huang CC, Deo RC. Phenomapping for novel classification of heart failure with preserved ejection fraction. Circulation. 2015;131:269–279. doi: 10.1161/CIRCULATIONAHA.114.010637

28. Peters AE, Tromp J, Shah SJ, Lam CSP, Lewis GD, Borlaug BA, Sharma K, Pandey A, Sweitzer NK, Kitzman DW, et al. Phenomapping in heart failure with preserved ejection fraction: insights, limitations, and future directions. Cardiovasc Res. 2023;118:3403–3415. doi: 10.1093/cvr/cvac179

29. Cohen JB, Schrauben SJ, Zhao L, Basso MD, Cvijic ME, Li Z, Yarde M, Wang Z, Bhattacharya PT, Chirinos DA, et al. Clinical Phenogroups in Heart Failure With Preserved Ejection Fraction: Detailed Phenotypes, Prognosis, and Response to Spironolactone. JACC Heart Fail. 2020;8:172–184. doi: 10.1016/j.jchf.2019.09.009

30. Hahn VS, Knutsdottir H, Luo X, Bedi K, Margulies KB, Haldar SM, Stolina M, Yin J, Khakoo AY, Vaishnav J, et al. Myocardial Gene Expression Signatures in Human Heart Failure With Preserved Ejection Fraction. Circulation. 2021;143:120–134. doi: 10.1161/CIRCULATIONAHA.120.050498

31. Hahn VS, Petucci C, Kim MS, Bedi KC, Jr., Wang H, Mishra S, Koleini N, Yoo EJ, Margulies KB, Arany Z, et al. Myocardial Metabolomics of Human Heart Failure With Preserved Ejection Fraction. Circulation. 2023;147:1147–1161. doi: 10.1161/CIRCULATIONAHA.122.061846

32. Pandey A, Parashar A, Kumbhani D, Agarwal S, Garg J, Kitzman D, Levine B, Drazner M, Berry J. Exercise training in patients with heart failure and preserved ejection fraction: meta-analysis of randomized control trials. Circ Heart Fail. 2015;8:33–40. doi: 10.1161/CIRCHEARTFAILURE.114.001615

33. Vacca A, Wang R, Nambiar N, Capone F, Farrelly C, Mostafa A, Sechi LA, Schiattarella GG. Lifestyle interventions in cardiometabolic HFpEF: dietary and exercise modalities. Heart Fail Rev. 2024. doi: 10.1007/s10741-024-10439-1

34. Lee VYJ, Houston L, Perkovic A, Barraclough JY, Sweeting A, Yu J, Fletcher RA, Arnott C. The Effect of Weight Loss Through Lifestyle Interventions in Patients With Heart Failure With Preserved Ejection Fraction-A Systematic Review and Meta-Analysis of Randomised Controlled Trials. Heart Lung Circ. 2024;33:197–208. doi: 10.1016/j.hlc.2023.11.022

35. Cuspidi C, Rescaldani M, Tadic M, Sala C, Grassi G. Effects of bariatric surgery on cardiac structure and function: a systematic review and meta-analysis. Am J Hypertens. 2014;27:146–156. doi: 10.1093/ajh/hpt215

36. Huang S, Lan Y, Zhang C, Zhang J, Zhou Z. The Early Effects of Bariatric Surgery on Cardiac Structure and Function: a Systematic Review and Meta-Analysis. Obes Surg. 2023;33:453–468. doi: 10.1007/s11695-022-06366-5

37. Sorimachi H, Obokata M, Omote K, Reddy YNV, Takahashi N, Koepp KE, Ng ACT, Rider OJ, Borlaug BA. Long-Term Changes in Cardiac Structure and Function Following Bariatric Surgery. J Am Coll Cardiol. 2022;80:1501–1512. doi: 10.1016/j.jacc.2022.08.738

38. Butler J, Shah SJ, Petrie MC, Borlaug BA, Abildstrom SZ, Davies MJ, Hovingh GK, Kitzman DW, Moller DV, Verma S, et al. Semaglutide versus placebo in people with obesity-related heart failure with preserved ejection fraction: a pooled analysis of the STEP-HFpEF and STEP-HFpEF DM randomised trials. Lancet. 2024;403:1635– 1648. doi: 10.1016/S0140-6736(24)00469-0

39. Tsujimoto T, Kajio H. Abdominal Obesity Is Associated With an Increased Risk of All-Cause Mortality in Patients With HFpEF. J Am Coll Cardiol. 2017;70:2739–2749. doi: 10.1016/j.jacc.2017.09.1111

40. Borlaug BA, Kitzman DW, Davies MJ, Rasmussen S, Barros E, Butler J, Einfeldt MN, Hovingh GK, Moller DV, Petrie MC, et al. Semaglutide in HFpEF across obesity class and by body weight reduction: a prespecified analysis of the STEP-HFpEF trial. Nat Med. 2023;29:2358–2365. doi: 10.1038/s41591-023-02526-x

41. Bagheri M, Agrawal V, Annis J, Shi M, Ferguson JF, Freiberg MS, Mosley JD, Brittain EL. Genetics of Pulmonary Pressure and Right Ventricle Stress Identify Diabetes as a Causal Risk Factor. J Am Heart Assoc. 2023;12:e029190. doi: 10.1161/JAHA.122.029190

42. Agrawal V, Manouchehri A, Vaitinadin NS, Shi M, Bagheri M, Gupta DK, Kullo IJ, Luo Y, McNally EM, Puckelwartz MJ, et al. Identification of Clinical Drivers of Left Atrial Enlargement Through Genomics of Left Atrial Size. Circ Heart Fail. 2024;17:e010557. doi: 10.1161/CIRCHEARTFAILURE.123.010557

43. Wei W, Riley NM, Lyu X, Shen X, Guo J, Raun SH, Zhao M, Moya-Garzon MD, Basu H, Sheng-Hwa Tung A, et al. Organism-wide, cell-type-specific secretome mapping of exercise training in mice. Cell Metab. 2023;35:1261–1279 e1211. doi: 10.1016/j.cmet.2023.04.011

44. Ramirez MF, Lau ES, Parekh JK, Pan AS, Owunna N, Wang D, McNeill JN, Malhotra R, Nayor M, Lewis GD, et al. Obesity-Related Biomarkers Are Associated With Exercise Intolerance and HFpEF. Circ Heart Fail. 2023;16:e010618. doi: 10.1161/CIRCHEARTFAILURE.123.010618

45. Fuster JJ, Zuriaga MA, Ngo DT, Farb MG, Aprahamian T, Yamaguchi TP, Gokce N, Walsh K. Noncanonical Wnt signaling promotes obesity-induced adipose tissue inflammation and metabolic dysfunction independent of adipose tissue expansion. Diabetes. 2015;64:1235–1248. doi: 10.2337/db14-1164

46. Tong S, Du Y, Ji Q, Dong R, Cao J, Wang Z, Li W, Zeng M, Chen H, Zhu X, et al. Expression of Sfrp5/Wnt5a in human epicardial adipose tissue and their relationship with coronary artery disease. Life Sci. 2020;245:117338. doi: 10.1016/j.lfs.2020.117338

47. Gorter TM, van Woerden G, Rienstra M, Dickinson MG, Hummel YM, Voors AA, Hoendermis ES, van Veldhuisen DJ. Epicardial Adipose Tissue and Invasive Hemodynamics in Heart Failure With Preserved Ejection Fraction. JACC Heart Fail. 2020;8:667–676. doi: 10.1016/j.jchf.2020.06.003

48. Crum Y, van Veldhuisen DJ, Shah SJ, Mylavarapu U, Hu M, Komtebedde J, Lobeek M, Hoendermis ES, Litwin SE, Solomon SD, et al. Epicardial adipose tissue, functional status and invasive hemodynamics in heart failure with preserved ejection fraction. JACC Adv. 2025:102185. doi: 10.1016/j.jacadv.2025.102185

49. Edwards JJ, Brandimarto J, Hu DQ, Jeong S, Yucel N, Li L, Bedi KC, Jr., Wada S, Murashige D, Hwang HTV, et al. Noncanonical WNT Activation in Human Right Ventricular Heart Failure. Front Cardiovasc Med. 2020;7:582407. doi: 10.3389/fcvm.2020.582407

50. Wu H, Tang LX, Wang XM, Li LP, Chen XK, He YJ, Yang DZ, Shi Y, Shou JL, Zhang ZS, et al. Porcupine inhibitor CGX1321 alleviates heart failure with preserved ejection fraction in mice by blocking WNT signaling. Acta Pharmacol Sin. 2023;44:1149– 1160. doi: 10.1038/s41401-022-01025-y

51. Paul MA, Wainwright CL, Hector EE, Ryberg E, Leslie SJ, Walsh SK. Short-Term Oral Administration of the Porcupine Inhibitor, Wnt-c59, Improves the Structural and Functional Features of Experimental HFpEF. Pharmacol Res Perspect. 2025;13:e70054. doi: 10.1002/prp2.70054

52. Wang TJ, Larson MG, Levy D, Benjamin EJ, Leip EP, Omland T, Wolf PA, Vasan RS. Plasma natriuretic peptide levels and the risk of cardiovascular events and death. N Engl J Med. 2004;350:655–663. doi: 10.1056/NEJMoa031994

53. McMurray JJ, Packer M, Desai AS, Gong J, Lefkowitz MP, Rizkala AR, Rouleau JL, Shi VC, Solomon SD, Swedberg K, et al. Angiotensin-neprilysin inhibition versus enalapril in heart failure. N Engl J Med. 2014;371:993–1004. doi: 10.1056/NEJMoa1409077

54. Bachmann KN, Gupta DK, Xu M, Brittain E, Farber-Eger E, Arora P, Collins S, Wells QS, Wang TJ. Unexpectedly Low Natriuretic Peptide Levels in Patients With Heart Failure. JACC Heart Fail. 2021;9:192–200. doi: 10.1016/j.jchf.2020.10.008

55. Lee DI, Zhu G, Sasaki T, Cho GS, Hamdani N, Holewinski R, Jo SH, Danner T, Zhang M, Rainer PP, et al. Phosphodiesterase 9A controls nitric-oxide-independent cGMP and hypertrophic heart disease. Nature. 2015;519:472–476. doi: 10.1038/nature14332

56. Cheng C, Zhang J, Li X, Xue F, Cao L, Meng L, Sui W, Zhang M, Zhao Y, Xi B, et al. NPRC deletion mitigated atherosclerosis by inhibiting oxidative stress, inflammation and apoptosis in ApoE knockout mice. Signal Transduct Target Ther. 2023;8:290. doi: 10.1038/s41392-023-01560-y

57. Wang X, Li J, Ma W, Meng L, Lu Y, Song J, Chen S, Zhen J, Yu X, Xi B, et al. Podocyte NPRC Deficiency Attenuates Glomerular Fibrosis in Diabetic Mice. Circ Res. 2025;137:e88–e105. doi: 10.1161/CIRCRESAHA.124.325702

58. Armstrong DWJ, Riley LA, Su YR, Shah AS, Absi T, Gupta DK, Wells QS, Brinkley DM, Stevenson LW, Merryman WD. Myocardial Neprilysin Is Increased in Hypertrophic Cardiomyopathy. Circulation. 2023;148:167–169. doi: 10.1161/CIRCULATIONAHA.123.064153

59. Packer M. The Adipokine Hypothesis of Heart Failure With a Preserved Ejection Fraction: A Novel Framework to Explain Pathogenesis and Guide Treatment. J Am Coll Cardiol. 2025;86:1269–1373. doi: 10.1016/j.jacc.2025.06.055

